# *Spherical*: an iterative workflow for assembling metagenomic datasets

**DOI:** 10.1101/067256

**Authors:** Thomas Hitch, Christopher J Creevey

**Author notes:** Contact information: Chris Creevey.

## Abstract

The consensus emerging from microbiome studies is that they are far more complex than previously thought, requiring deep sequencing. As deep sequenced datasets provide greater coverage than previous datasets, recovering a higher proportion of reads to the assembly is still a challenge. To tackle this issue, we set of to identify if multiple iterations of assembly would allow for otherwise lost contigs to be formed and studied and if so, how successful is such an avenue at improving the current methodology.

A simulated metagenomic dataset was initially used to identify if multiple iterations of assembly produce useable contigs or mis-assembled artefacts were produced. Once we had confirmed that the secondary iterations were producing both accurate contigs without a reduction in contig quality we applied this methodology in the form of *Spherical* to 3 metagenomic studies.

The additional contigs produced by Spherical increased the number of reads aligning to an identified gene by 11–109% compared to the initial iterations assembly. As the size of the dataset increased, as did the amount of data multiple iterations were able to add.

**Availability:** Spherical is implemented in Python 2.7 and available for use under a MIT licence agreement at: https://github.com/thh32/Spherical

## Introduction

Over the last 10 years researchers have utilised high-throughput sequencing to investigate the structure and function of increasing larger numbers of microbial communities from environments across the globe (Krohn-Molt et al. 2013; van der Lelie et al. 2012; Modi et al. 2013). While these studies have provided unique and novel insights into the workings of these communities, there is a growing consensus that the tools available are not capturing the full functional diversity of the data being generated (Nagarajan & Pop 2013). Increasing the proportion of genome assembled is a challenge, and resolving this issue is very needed, with the ever increasing amounts of sequencing data becoming available. Additionally, it would also enable us to retrieve further more from data already available in repositories.

Mathematically, *de novo* assembly of a genome falls within the class of problems for which no efficient algorithm is known (NP-hard) (Myers. 1995; Medvedev et al. 2007), leading to the proposal of a variety of heuristic solutions (Charuvaka & Rangwala 2011). These have ranged from simple overlap-layout-consensus approaches, where sequencing reads with overlapping regions are joined together into contigs (Myers. 1995) to more complex approaches such as de Bruijn graphs (Idury et al. 1995). Genomic assemblers based on de Bruijn graphs break each read into smaller strings of K length (kmers) and connections are made between overlapping kmers (Schatz et al. 2010). Paths of least resistance through the kmer graph represent contigs (Compeau et al. 2011).

Generally, these assemblers have been designed with a single-genome in mind where it could be guaranteed that all sequences generated belonged to the same species. However, as sequencing approaches for sampling the genomic information of entire microbial communities (metagenomics) began to emerge (Venter et al. 2004), it was clear that new approaches may be necessary (Lai et al. 2012). By far the biggest issue with metagenomic datasets stem from the uneven distribution of species in natural communities leading to varying depths of sequencing coverage of each (Nagarajan & Pop 2013). This leads to over-sequencing of dominant species in the community and results in heavily fragmented assemblies of the genomes of minority species, if they can be assembled at all (Bergeron et al. 2007).

Promising solutions to dealing with this problem use a ‘divide and conquer’ approach to break the data into more easily manageable pieces (Bergeron et al. 2007). For example the data from environmental samples can be split into “bins” representing different taxa from the community (Mohammed et al. 2011). Sequence reads can be sorted into bins based on properties such as kmer-frequency or the percentage of Guanine and Cytosines (GC) they contain (Dro & Mchardy 2012). This has the potential to increase the assembly rate of low abundance species, however it depends heavily on accurately partitioning the data (Wang et al. 2012). Indeed, bins of metagenomic data produced in this manner may represent a single species or an entire phylum depending on the complexity of the community (Wang et al. 2012).

Digital normalization is another method that tries to deal with unevenly sampled data (Brown & Crusoe 2014). In this approach, sequence reads are discarded based on the local coverage of their kmers (Brown & Crusoe 2014). This has the effect of reducing coverage of over-represented taxa, and “normalising” the coverage to make it more even (Brown & Crusoe 2014). While this pre-processing step allows for reduction of the datasets size, it does not reduce the complexity of metagenomes to be assembled.

Using a similar approach to ‘divide-and-conquer’ large genomic datasets, SLICEMBLER aims to improve deep sequenced (>800x coverage) single genome assemblies (Mirebrahim et al. 2015). The approach taken here is to separate the input data into pre-determined “slices” based on coverage (Mirebrahim et al. 2015). Each slice is then assembled separately and frequently occurring strings (FOS) are identified between the sub-assemblies and merged, effectively scaffolding them together to produce a final assembly (Mirebrahim et al. 2015). This approach works well for deep sequenced genomic datasets where coverage is known, however in metagenomics datasets from uncharacterised microbial communities, coverage is generally unknown (Peng et al. 2012).

Whilst these approaches are an improvement upon single-genome based methods, they still do not generate an assembly that utilises 100% of the reads from the data or produce complete genomes without manual curation (Hess et al. 2011).

Here we propose an iterative workflow that tries to address this problem with metagenomic assemblies. Based on, and extending the ‘divide-and-conquer’ principle, the workflow (called “Spherical”) can be applied to any assembly method. Spherical is designed to work with datasets that cannot be assembled in a single step due to data volume or complexity but could equally be applied to smaller and simpler datasets.

## *Spherical* workflow

Spherical uses an iterative workflow which tries to capture the reads not utilised by an assembly, and these are reapplied to subsequent rounds of assemblies. The outline of the approach is summarised in Figure 1.

**Figure 1:**
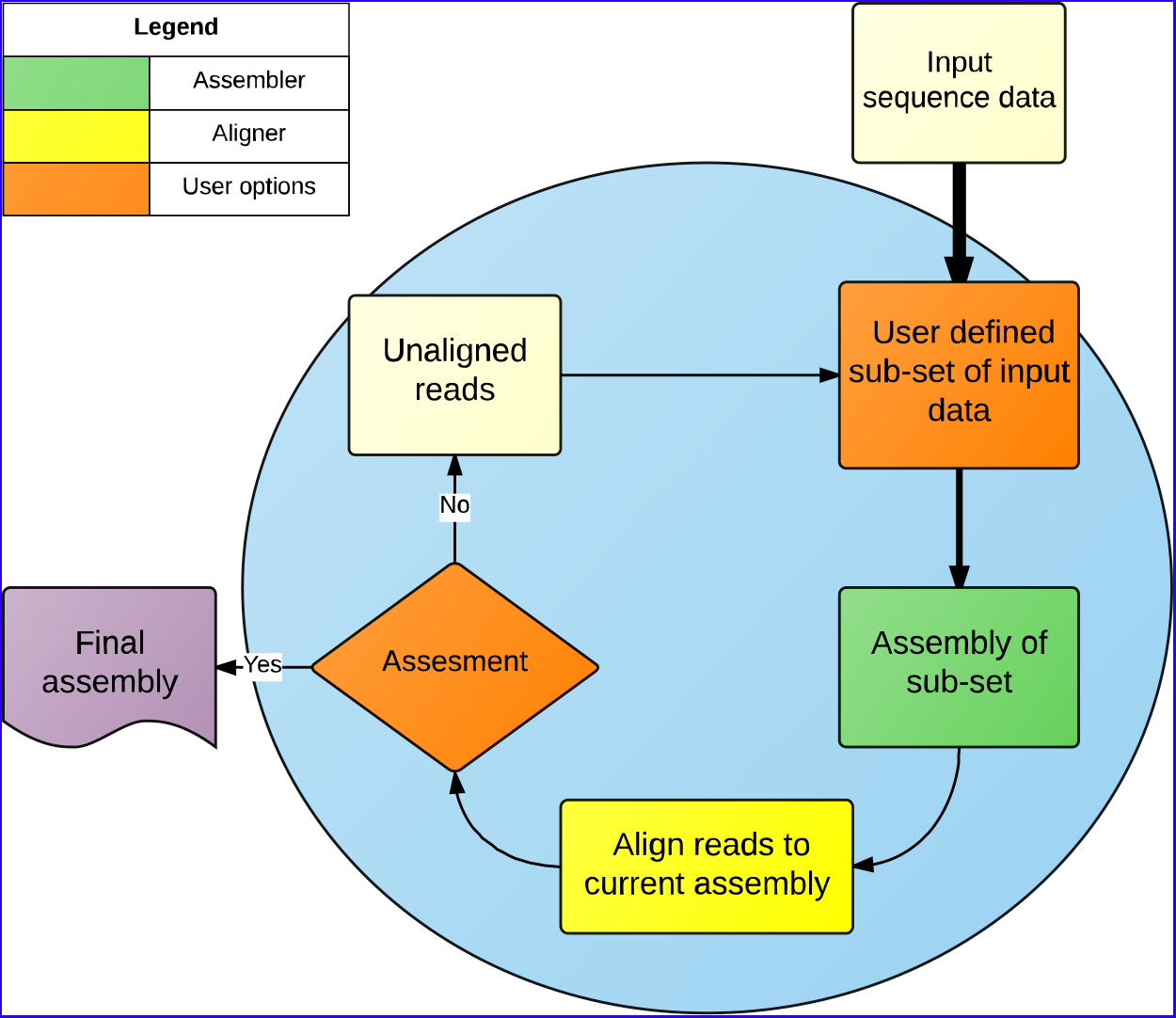
A flow-chart of the steps used by *Spherical*. The all process within the blue circle occur within Spherical whilst the input and output files are outside. The width of the arrows indicate the possible decrease in file size depending on user sub-sample selection. The ‘user defined criteria’ is defined as any user option which indicates a point at which Spherical should stop iterating.

The Spherical workflow is composed of 5 steps:

### Step 1: Sub-sample selection

The first step in *Spherical* is the optional initial sub-sampling of the sequencing data. This can be advantageous when the working with very large datasets. In this process a random sub-sample (defined by –R) is taken from the input sequencing data. Using a subset fraction of 1 selects the entire input dataset instead. If only one value is given to –R, then Sphercial will apply this sub-sample fraction at every iteration, however the user also has the option of providing values to be used at each iteration.

### Step 2: Assembly

The sub-sample is then assembled using the assembler of choice. Currently the default assembler in Spherical is Velvet (Namiki et al. 2012).

### Step 3: Alignment

When the assembly is completed Spherical uses Bowtie 2 (Langmead & Salzberg 2012) to align all reads previously unaligned (in iteration 1 this is all the reads) to the contigs resulting from this assembly. All reads that do not align to the assembly produced at this iteration are saved for the subsequent rounds. If a read aligns to the assembly at this point, it is considered utilised and hence excluded from the subsequent iterations.

### Step 4: Assessment

The user can define two parameters to be used by Spherical to determine completeness. The first is based on the number of iterations currently completed (iter). When spherical has completed all iterations defined by this value, it will halt carrying out iterations, and move to step 5. The second option is based on the proportion of reads currently utilised by the total assembly (-align). This is calculated as the number of reads currently unaligned, divided by the total number of reads initially provided. When Spherical determines that the alignment rate has been reached it will halt carrying out iterations and move to step 5. The user must provide these two parameters, the current default is 5 iterations or an alignment rate of 70%.

If neither of these criteria have been met, spherical will pass on all unaligned reads to step 1 for another iteration.

### Step 5: Final output

When Spherical has met the user-defined criteria for halting, Spherical will combine all the assemblies from each iteration into a single file and calculates statistics such as N50, lengths of longest and shortest contigs the standard deviation of the lengths and the alignment rate both for each iteration and for the final assembly.

In the following sections we outline the principles behind Spherical and demonstrate its use on three different metagenomic datasets of differing sizes and complexity. We show that by taking an iterative approach the resulting assemblies use a greater proportion of the original raw reads and in large datasets it allowed to retrieve more information from the less-represented organisms in the community.

## Results

### Simulated dataset analysis

By using a simulated dataset (Mende et al. 2012) created from 400 species of varying abundance we investigated the accuracy of assemblies from each iteration from the Spherical workflow. We used BLASTN to identify the best matching genomic region for each contig assembled. The quality of the reconstructed region was assessed using the contig score (Mende et al. 2012) which calculates a value representing the accuracy of the reconstruction. We found that there was no observable decrease in the contig score as the number of iterations increased (Supplementary Table 3).

### Metagenome dataset analysis

We used three published metagenomic datasets (chicken ceacum (Qu et al. 2008), human oral (Belda-Ferre et al. 2012) and groundwater from the Yucatan peninsula (Moore et al. 2009)) of varying sizes to test Spherical (Table 2). For each we carried out three assembly approaches, 1) Basic Velvet assembly, 2) Digital normalisation followed by a Velvet assembly and 3) Spherical on the raw data with 5 iterations, using Velvet as the assembler (Supplementary Table 1). We used the percentage of the assembly to which no reads align as a measure of miss-assembly (the “false base rate”) and the percentage of reads that aligned as a measure of completeness.

**Table 1:** Number of reads assigned to an identified gene in each Spherical assembly.

| Dataset | Iteration 1 | Iteration 2 | Iteration 3 | Iteration 4 | Iteration 5 |
| --- | --- | --- | --- | --- | --- |
| Cecum | 164,266 | 11,317 | 3,398 | 2,309 | 1,427 |
| Oral | 126,388 | 43,721 | 33,110 | 30,907 | 30,032 |
| Groundwater | 130,451,683 | 27,463,983 | 12,121,835 | 8,586,974 | 7,783,534 |

**Table 2:**
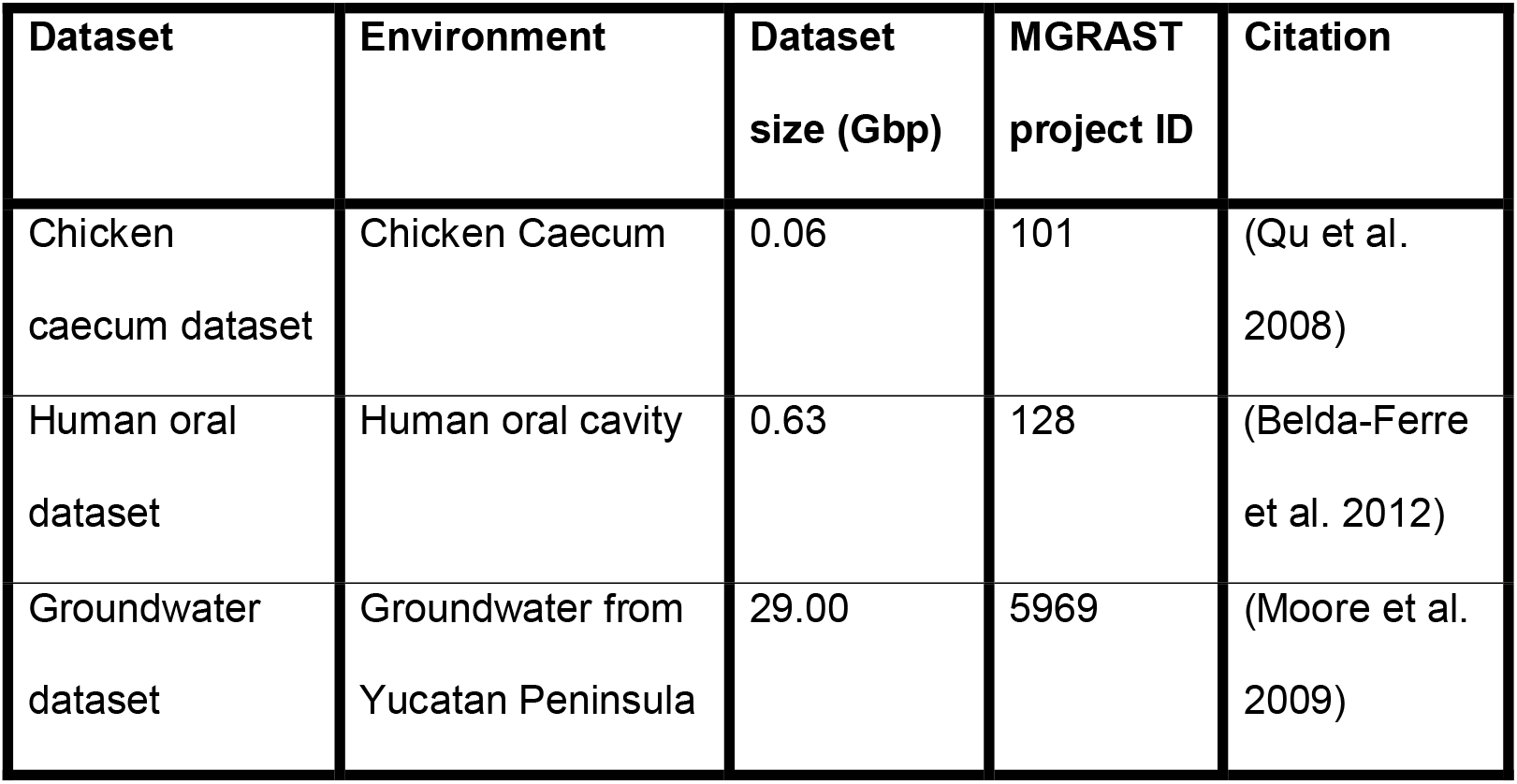
Information on each dataset

| Dataset | Environment | Dataset size (Gbp) | MGRAST project ID | Citation |
| --- | --- | --- | --- | --- |
| Chicken caecum dataset | Chicken Caecum | 0.06 | 101 | (Qu et al. 2008) |
| Human oral dataset | Human oral cavity | 0.63 | 128 | (Belda-Ferre et al. 2012) |
| Groundwater dataset | Groundwater from Yucatan Peninsula | 29.00 | 5969 | (Moore et al. 2009) |

### Assembly quality for tested datasets

The Chicken Caecum microbiome was the smallest of the three datasets tested. As a result, all three assembly approaches produced very similar results (Supplementary Table 1). However the assembly from Spherical utilised 1% more of the raw reads than the other approaches. This was at the cost of slightly lowering the N50 (from 109 to 104) and increased false base rate (from 0.01% to 0.04%).

The human oral dataset was a larger dataset, and as a result we observed a greater variability in how the different assembly approaches performed. Spherical was able to increase both the N50 (from 190 to 234) and the alignment rate (from 13% to 24.6%) compared to the next best approach (basic velvet assembly), however the false base rate also increased (from 0.02% to 0.19%).

The groundwater dataset was the largest tested. The alignment rate of Spherical increased (from 52.8% to 59.7%) and false base rate decreased (from 3.86% to 2.89%), however *Sphericals* N50 was significantly reduced (from 330 to 211) in comparison to the normalised assembly.

### Effect of sub-sample size on the resulting assembly

To study the effect of altering the sub-sample size we used the largest of the metagenomics datasets tested (Groundwater) and with sub-sample fractions of 1 (representing 1/1, i.e. all the raw data), 4 (representing ¼ of the raw sequencing reads) and 30 (representing 1/30 of the raw sequencing reads). As shown in Supplementary Table 1 the change in sub-sample size resulted in a small change in the quality of the resulting assembly; increasing the false base rate from 2.89% to 3.78%, and reducing the N50 from 211 to 189 and reducing the alignment rate from 59.7% to 49.8%. However even with these changes, the taxonomic profile of the each assembly did not differ, Supplementary Figure 1.

### Effect of multiple iterations of assembly

The additional iterations employed by Spherical lead to an increase in the number of reads that could be assigned to known genes (Figures 3,4 & 5). As shown in Figure 3 and 4, for small metagenomic datasets the taxonomic profile does not change across iterations, however the iterative approach does allow for almost twice the number of reads to be assigned to a taxonomic class (Table 1). In the groundwater dataset the secondary iterations provided a different taxonomic profile compared to the initial iteration (Figure 5).

**Figure 3:**
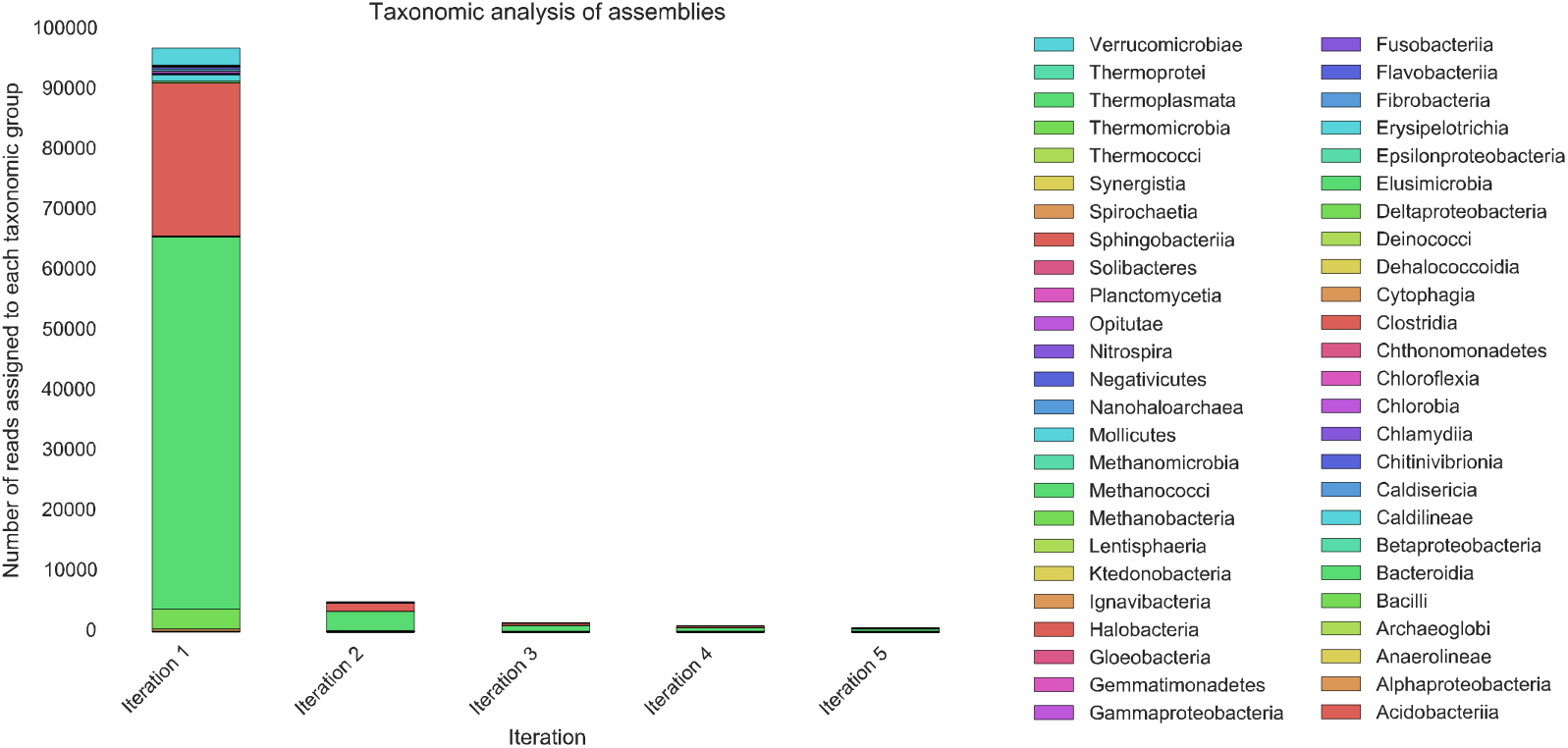
A taxonomic breakdown of each iteration of cecum dataset Spherical assembly at the class level.

**Figure 4:**
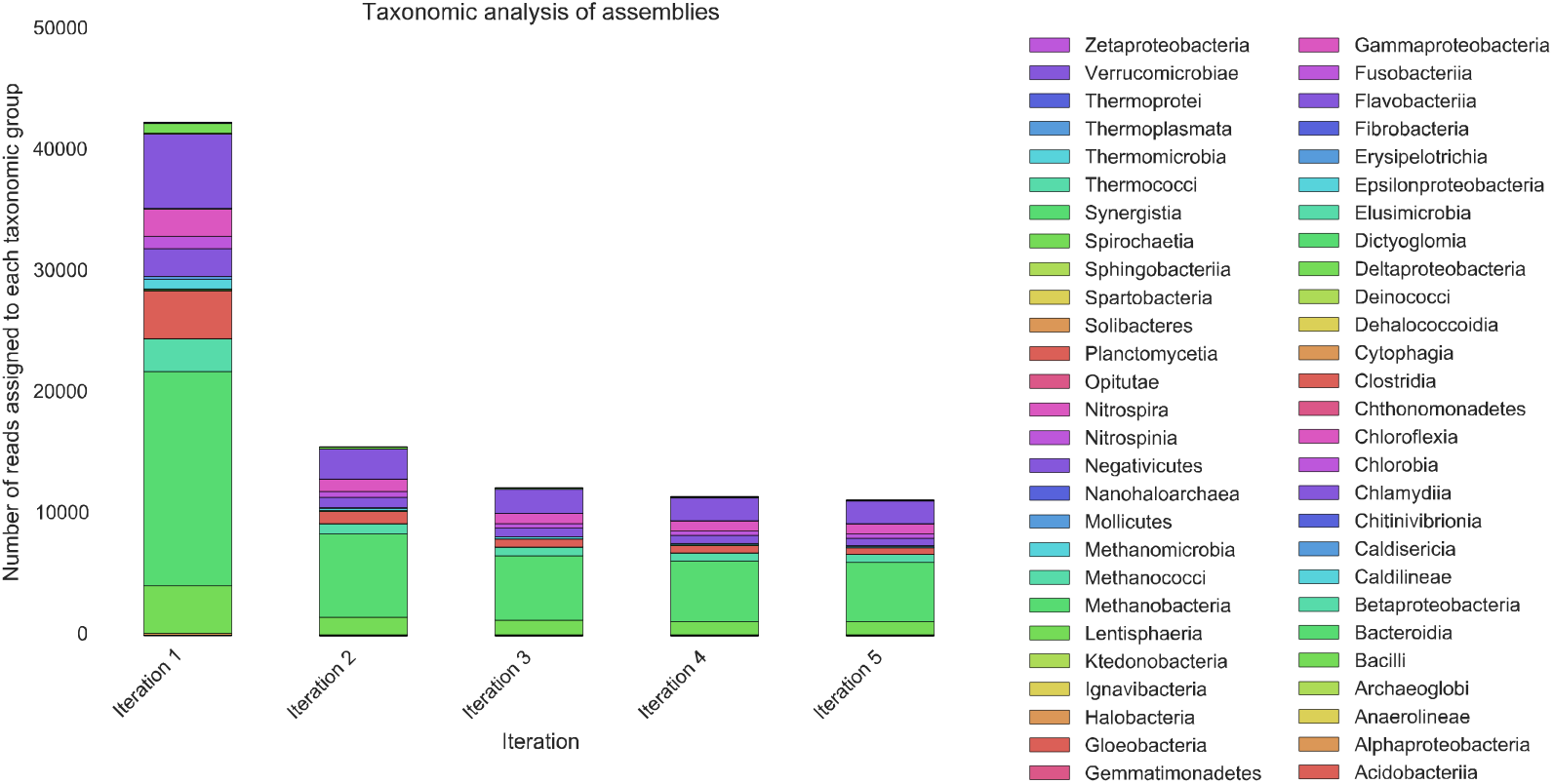
A taxonomic breakdown of each iteration of oral dataset Spherical assembly at the class level.

**Figure 5:**
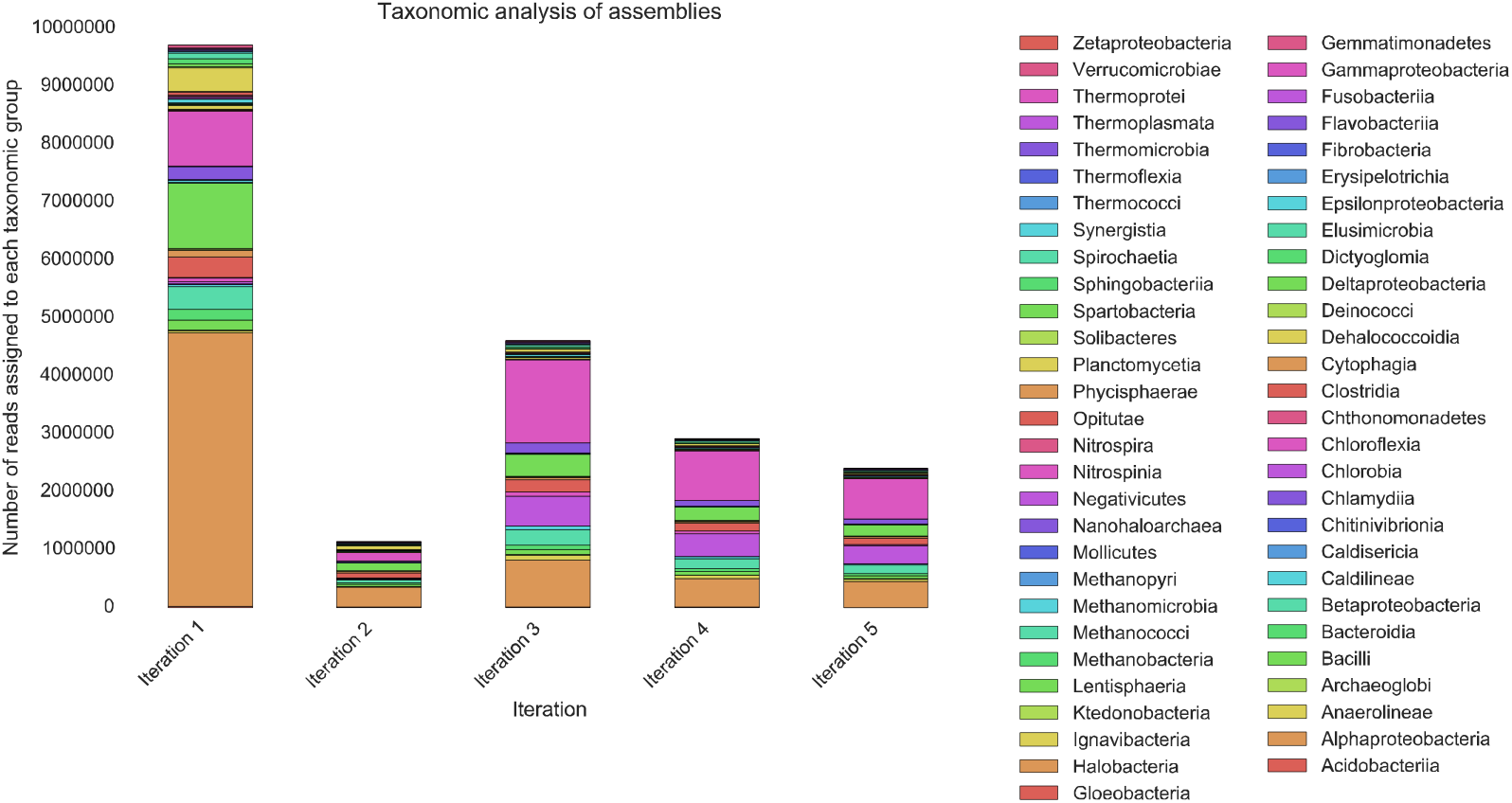
A taxonomic breakdown of each iteration of groundwater dataset *Spherical* (subset size = 1) assembly at the class level.

As shown in Table 1, the additional iterations allowed for identification of 11% additional reads in the chicken caecum dataset, 109% additional reads in the human oral dataset and 43% additional reads in the groundwater dataset compared to the first iterations assembly.

## Methods

### Simulated dataset

A simulated Illumina sequenced metagenome of 400 species, each species abundance within the dataset differed to produce a more accurate representation of a metagenomic dataset (Mende et al. 2012). The raw data was downloaded and assembled using a kmer of 31 and subset size of 1 with Spherical. Contig scores were calculated for each contig produced using the method described in Mende et al, 2012.

### Datasets

We have selected 3 metagenomic samples from different environments to allow for comparison of the methods on datasets with different sequencing depths and complexity, Table 2.

All of the datasets used were obtained from MG-RAST (Meyer et al. 2008), Table 2, and aimed to provide variation in sampling environments as to provide a robust testing sample for *Spherical*.

### Methods of assembly

We chose digital normalisation as comparative method to *Spherical* due to its ability to produce a subset of the dataset with uniform coverage without removing the datasets complexity. SLICEMBLER was also considered however as covered earlier metagenomics provides an unknown coverage due to the microbial population itself being unknown and therefore cannot be supplied to SLICEMBLER.

The optimum kmer size was identified before the comparison assemblies by assembly of each dataset using Velvet with a kmer of 21,31,41 and 51. The raw reads were then aligned to each assembly using Bowtie2. The assembly with the highest alignment rate was then selected and that kmer used for the three methods being compared. All the methods used Velvet as the assembler, removing the assembler as a variable. Velvet was selected as the base assembler due to being open-source, allowing anyone access to it, as well as producing high quality metagenomic assemblies (Shi et al. 2014; Namiki et al. 2012).

### Khmer - Digital normalization

Digital normalization is based on the use of kmers and is part of the Khmer package. Firstly Khmer breaks each read into kmers, producing a hash-table for the entire dataset present in the dataset. Once counted, it removes reads consisting of redundant kmers, reducing the dataset size whilst keeping sufficient data for an accurate assembly. With each dataset we applied the “normalize-by-median.py” script of Khmer, with a kmer of 20, 4 hash tables of size 32e9 and an ideal median of 20.

### Spherical

*Spherical* was run on each dataset with variable sub-set sizes, ranging from 1 to 30. This allowed us to explore the effect of the sub-sampling function of *Spherical*.

### Basic assembly method

The basic assembly method involved the entire dataset being entered into Velvet with the optimum kmer and the expected coverage is automatically selected by Velvet.

### Assembly quality

As we are unable to produce contig scores for real metagenomic assemblies we used the following statistics to provide insight in the quality of each assembly; alignment rate, N50 and false base rate. The alignment rate identifies how much of the raw data is accounted for by the assembly. The higher the alignment rate the more representative the assembly is of the entire dataset. N50 identifies how fragmented the assembly is by identifying average contig length. False bases (bases to which no read aligns) have no basis in the dataset and therefore indicate the rate of misassembled contigs within an assembly.

### RAM usage

*Spherical* has been designed to reduce RAM usage, in the hope of overcoming the issue of limited computing infrastructure. When comparing methods, maximum RAM used was taken into account. For normalisation, RAM usage was monitored during both the normalisation and assembly stages, the peak was then taken as max RAM usage. The basic assembly methods RAM usage was also monitored to be used as a base line for RAM requirement for assembly of the dataset.

### Taxonomic analysis

Whilst statistics about the assembly allow us to identify how representative the assembly is of the raw data, taxonomic annotation allows us greater insight into the data. Each assembly was taxonomically annotated against the bacterial and archaeal UNIPROT databases using RAPsearch (Zhao et al. 2012; Consortium 2015). The output was then converted into a general feature format (GFF) file and filtered by overlap using MGKIT. HTSeq-count was then used to identify the abundance of phylums within each assembly (Anders et al. 2015).

### Effect of *Spherical* subset function

To study the effect *Spherical* sub-sampling had on the output, dataset 3 was assembled under 3 different sub-sampling levels. Firstly, the entire file was included (sub-sampling = 1). This allowed us to study what the basic effect of multiple iterations of assembly was on the output. Next, a quarter of all reads were used (subsampling = 4), allowing us to study the effect whilst significantly reducing RAM usage (but still aiming to produce a quality assembly). Finally, only one thirtieth of the reads were used (set-sampling = 30), this was aimed to study the extreme effects of subsampling a very small amount of reads and how this would effect the output assembly as well as RAM usage.

### Effect of multiple iterations of assembly

The *Spherical* assembly of the Cecum, Oral and Groundwater datasets (subsampling = 1) were studied to understand the biodiversity within each iterations of assembly. The ability of multiple iterations to uncover low abundant species was also studied by taxonomically annotating (against bacterial and archaeal UNIPROT database using RAPsearch) each iteration and evaluating their taxonomic profile.

## Acknowledgments

We wish to thank the advice provided by Francesco Rubino to this project. TH was funded through the New Zealand fund for Global Partnerships in Livestock Emissions Research (GPLER). CJC was funded under the Biotechnology and Biological Sciences Research Council (BBSRC) Institute Strategic Programme Grant, Rumen Systems Biology, (BB/E/W/10964A01).

## References

Anders, S., Pyl, P.T. & Huber, W., 2015. Genome analysis HTSeq — a Python framework to work with high-throughput sequencing data., 31(2), pp.166–169.

Belda-Ferre, P. et al., 2012. The oral metagenome in health and disease. The ISME Journal, 6(1), pp.46–56. Available at: http://www.nature.com/doifinder/10.1038/ismej.2011.85.

Bergeron, A. et al., 2007. Divide and Conquer□: Enriching Environmental Sequencing Data., (9).

Charuvaka, A. & Rangwala, H., 2011. Evaluation of short read metagenomic assembly. BMC genomics, 12 Suppl 2(Suppl 2), p.S8. Available at:http://www.pubmedcentral.nih.gov/articlerender.fcgi?artid=3194239&tool=pmcentrez&rendertype=abstract [Accessed January 21, 2014].

Compeau, P.E.C., Pevzner, P. a & Tesler, G., 2011. How to apply de Bruijn graphs to genome assembly. Nature biotechnology, 29(11), pp.987–91. Available at: http://www.ncbi.nlm.nih.gov/pubmed/22068540 [Accessed July 9, 2014].

Consortium, T.U., 2015. UniProt□: a hub for protein information., 43(October 2014), pp.204–212.

Dro, J. & Mchardy, A.C., 2012. axonomic binning of metagenome samples generated by next-generation sequencing technologies., 13(6), pp.646–655.

Hess, M. et al., 2011. Metagenomic discovery of biomass-degrading genes and genomes from cow rumen. Science (New York, N.Y.), 331(6016), pp.463–7.

Krohn-Molt, I. et al., 2013. Metagenome survey of a multispecies and alga-associated biofilm revealed key elements of bacterial-algal interactions in photobioreactors. Applied and environmental microbiology, 79(20), pp.6196–206. Available at: http://www.ncbi.nlm.nih.gov/pubmed/23913425 [Accessed February 14, 2014].

Lai, B. et al., 2012. A de novo metagenomic assembly program for shotgun DNA reads. Bioinformatics (Oxford, England), 28(11), pp.1455–62. Available at: http://www.ncbi.nlm.nih.gov/pubmed/22495746 [Accessed January 23, 2014].

Langmead, B. & Salzberg, S.L., 2012. Fast gapped-read alignment with Bowtie 2. Nature Methods, 9(4), pp.357–359.

van der Lelie, D. et al., 2012. The metagenome of an anaerobic microbial community decomposing poplar wood chips. PloS one, 7(5), p.e36740. Available at: http://www.pubmedcentral.nih.gov/articlerender.fcgi?artid=3357426&tool=pmcentrez&rendertype=abstract [Accessed February 11, 2014].

Mende, D.R. et al., 2012. Assessment of metagenomic assembly using simulated next generation sequencing data. PloS one, 7(2), p.e31386. Available at: http://www.pubmedcentral.nih.gov/articlerender.fcgi?artid=3285633&tool=pmcentrez&rendertype=abstract [Accessed January 28, 2014].

Meyer, F. et al., 2008. The metagenomics RAST server – a public resource for the automatic phylogenetic and functional analysis of metagenomes., 8, pp.1–8.

Mirebrahim, H., Close, T.J. & Lonardi, S., 2015. De novo meta-assembly of ultradeep sequencing data. Bioinformatics, 31(12), pp.i9–i16. Available at: http://bioinformatics.oxfordjournals.org/content/31/12/i9.full?etoc.

Modi, S.R. et al., 2013. Antibiotic treatment expands the resistance reservoir and ecological network of the phage metagenome. Nature, 499(7457), pp.219–22. Available at: http://www.pubmedcentral.nih.gov/articlerender.fcgi?artid=3710538&tool=pmcentrez&rendertype=abstract [Accessed January 20, 2014].

Mohammed, M.H. et al., 2011. SPHINX--an algorithm for taxonomic binning of metagenomic sequences. Bioinformatics (Oxford, England), 27(1), pp.22–30. Available at: http://bioinformatics.oxfordjournals.org/cgi/content/long/27/1/22 [Accessed February 19, 2014].

Nagarajan, N. & Pop, M., 2013. Sequence assembly demystified. Nature reviews. Genetics, 14(3), pp.157–67. Available at: http://www.ncbi.nlm.nih.gov/pubmed/23358380 [Accessed January 21, 2014].

Namiki, T. et al., 2012. MetaVelvet: an extension of Velvet assembler to de novo metagenome assembly from short sequence reads. Nucleic acids research, 40(20), p.e155. Available at: http://www.pubmedcentral.nih.gov/articlerender.fcgi?artid=3488206&tool=pmcentrez&rendertype=abstract [Accessed May 26, 2014].

Peng, Y. et al., 2012. IDBA-UD: a de novo assembler for single-cell and metagenomic sequencing data with highly uneven depth. Bioinformatics (Oxford, England), 28(11), pp.1420–8. Available at: http://www.ncbi.nlm.nih.gov/pubmed/22495754 [Accessed February 4, 2014].

Qu, a et al., 2008. Comparative Metagenomics Reveals Host Specific Metavirulomes and Horizontal Gene Transfer Elements in the Chicken Cecum Microbiome. Plos One, 3(8), p.19. Available at: <Go to ISI>://WOS:000264412600031.

Shi, W. et al., 2014. Methane yield phenotypes linked to differential gene expression in the sheep rumen microbiome. Genome research. Available at: http://www.ncbi.nlm.nih.gov/pubmed/24907284 [Accessed June 10, 2014].

Venter, J.C. et al., 2004. Environmental genome shotgun sequencing of the Sargasso Sea. Science (New York, N.Y.), 304(5667), pp.66–74.

Wang, Y. et al., 2012. MetaCluster 5.0: a two-round binning approach for metagenomic data for low-abundance species in a noisy sample. Bioinformatics (Oxford, England), 28(18), pp.i356–i362. Available at: http://www.pubmedcentral.nih.gov/articlerender.fcgi?artid=3436824&tool=pmcentrez&rendertype=abstract [Accessed February 14, 2014].

Zhao, Y., Tang, H. & Ye, Y., 2012. RAPSearch2: a fast and memory-efficient protein similarity search tool for next-generation sequencing data. Bioinformatics (Oxford, England), 28(1), pp.125–6. Available at: http://www.pubmedcentral.nih.gov/articlerender.fcgi?artid=3244761&tool=pmcentrez&rendertype=abstract [Accessed January 25, 2015].

